# Evaluating Long-Range Temporal Structure in Foundation Model-Based Forecasts of Heartbeat Dynamics

**DOI:** 10.64898/2026.05.25.727760

**Authors:** Adrian Serapio, Bharath Ramsundar, Sandya Subramanian

## Abstract

We examine the long-range temporal structure of forecasts produced by Time-Series Foundation Models (TSFMs) on heartbeat dynamics using the MIT-BIH Normal Sinus Rhythm Database (NSRDB). Our findings indicate that these models do not adequately capture long-range dependencies, as reflected in growing errors in RR-interval predictions over longer forecast horizons. Code is available at https://github.com/SubramanianLab/ecg-tsfm-benchmark.

## 1. Introduction

Time-Series Foundation Models (TSFMs) have demonstrated remarkable capabilities in forecasting diverse temporal data (Liang et al., 2024). By leveraging large-scale pre-training on various time-series datasets, TSFMs are capable of forecasting without task-specific training. This in-context learning capability offers a promising solution to regimes where there is a scarcity of available data, particularly in the healthcare domain, where high-quality longitudinal data is often difficult to acquire.

Although TSFMs demonstrate the potential to greatly advance physiological modeling, it remains unclear whether such models are actually capable of modeling the long-range structure of physiological data. Particularly, we chose to focus on evaluating heartbeat dynamics and investigate whether TSFMs merely reproduce local beat-to-beat patterns or capture physiologically plausible heart rate dynamics over longer time scales.

We evaluate the ability of TSFMs to model the long-range structure of heartbeat dynamics by forecasting Electrocardiogram (ECG) data. First, ECG is a very wide-spread modality given its importance in cardiac monitoring (Siontis et al., 2021). Long-range ECG forecasting enables longitudinal adverse cardiac event prediction including earlier detection of sudden cardiac death, ventricular arrhythmias, and myocardial infarction which can greatly reduce cardiac disease-related morbidity and improve survival outcomes (Hernesniemi et al., 2026; DiMarco & Philbrick, 1990; Siontis et al., 2021).

Second, ECG data is characterized by a complex multi-scale temporal hierarchy, where fine-grained morphological features are nested within broader physiological trends (Draghici & Taylor, 2016). The focus of our work goes beyond the local waveform morphology (milliseconds) and more to the long-range RR interval structure of ECG data (minutes to hours). Measuring TSFMs’ ability to model RR interval structure is critical, as autonomic regulation signals emerge over longer time scales. RR interval forecasting provides a more direct probe of autonomic regulation and the body’s physiological control system.

Although ECG-specific foundation models have been proposed (Li et al., 2025a;b; Tang et al., 2025), the lack of publicly available weights and their predominant focus on classification rather than forecasting led us to focus our analysis on general-purpose time-series foundation models.

Motivated by the above, we made the following contributions:

1. **Long-Range Physiological Signal Evaluation**: Our analysis focuses on evaluating the capacity of TSFMs to forecast RR intervals, representing long-range temporal structure in the minutes to hours range than simply local waveform-level (milliseconds)
2. **Physiological Distributional-Level Evaluation**: We evaluate TSFMs beyond the typical metrics used typically limited to MAE and RMSE. We also include Standard Deviation of Normal-to-Normal Intervals (SDNN) and Root Mean Square of Successive Differences (RMSSD) in order to evaluate if the TSFMs evaluate the fluctuations of the RR interval signal. Moreover, we also include a Kolmogorov-Smirnov (KS) goodness-of-fit metric as a metric to measure alignment with the underlying statistical generative process governing heartbeat dynamics (Subramanian & Ramsundar, 2025).

Our findings underscore the necessity of evaluating TSFMs through the lens of physiological validity before they can be reliably deployed for clinical forecasting.

## 2. Materials and Methods

### 2.1. Dataset

We use the MIT-BIH normal sinus rhythm dataset for bench-marking the TSFMs (The Beth Israel Deaconess Medical Center, 1990). This dataset contains 18 ECG recordings of subjects referred to the Arrhythmia Laboratory at Boston’s Beth Israel Hospital (now the Beth Israel Deaconess Medical Center).

### 2.2. Time-Series Foundation Models

Although several foundation models have been developed for ECG and PPG data (Li et al., 2025a;b; Tang et al., 2025), most focus on classification or event detection rather than forecasting, and many lack publicly available weights, precluding inclusion in this study. While specialized time-series models exist (Nie et al., 2023), we instead evaluate the zero-shot forecasting capabilities of general-purpose Time-Series Foundation Models (TSFMs). This is motivated by the fact that modern TSFMs already incorporate patch-based transformer architectures (e.g., PatchTST), enabling modeling of long-range temporal dependencies. For comparison, we additionally include an LSTM as a non-foundation model baseline to contextualize the performance of the TSFMs.

We evaluate the following TSFMs (Table 1):

**Table 1.** Characteristics of Time-Series Foundation Models Bench-marked.

|  | TimesFM-2.5 | Chronos-2 | Moirai 2.0 |
| --- | --- | --- | --- |
| Number of Parameters | 200M | 120M | 11.4M |
| Context Length | 16,000 | 8,192 | — |
| Maximum Pred. Length | — | 1,024 | — |

1. **TimesFM 2.5 (Google):** A decoder-only transformer using patching to efficiently capture long-range dependencies, pre-trained on large-scale synthetic and real-world time-series data (Das et al., 2024).
2. **Chronos-2 (Amazon):** Frames forecasting as a language modeling problem by quantizing continuous values into discrete tokens and predicting their distribution autoregressively (Ansari et al., 2025).
3. **Moirai 2.0 (Salesforce):** A lightweight decoder-only TSFM with single-patch input representation and quantile loss for probabilistic forecasting (Liu et al., 2026).

### 2.3. Evaluation Metrics

To assess the accuracy and distributional fidelity of the forecasted ECG signals, we employed several primary metrics. Let *y*_*t*_ denote the ground-truth signal and *ŷ*_*t*_ the corresponding model prediction at time index *t*, where *t* = 1, …, *H* and *H* is the forecast horizon (number of predicted time points).

- **Mean Absolute Error (MAE):** Measures the average absolute magnitude of the forecast errors.

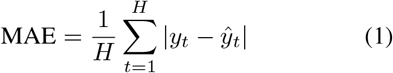
- **Root Mean Squared Error (RMSE):** Provides a metric that disproportionately penalizes large outliers, which is critical for identifying significant deviations in cardiac cycle timing.

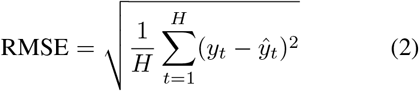
- **Standard Deviation of NN intervals (SDNN):** Measures the overall variability of the RR intervals (normal-to-normal intervals), reflecting both short- and longterm components of heart rate variability.

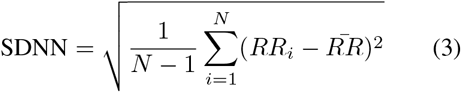

where *RR*_*i*_ denotes the *i*-th RR interval and 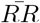 is the mean RR interval. SDNN captures global variability and is sensitive to long-range temporal structure in the heartbeat dynamics.
- **Root Mean Square of Successive Differences (RMSSD):** Quantifies short-term variability by measuring the root mean square of differences between successive RR intervals.

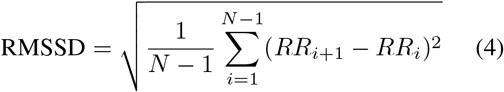 RMSSD is primarily sensitive to high-frequency variations and reflects parasympathetic (vagal) activity, making it useful for assessing short-term temporal fidelity of the forecasted signals.
- **Kolmogorov-Smirnov (KS) Score:** We use the KS score as a Goodness-of-Fit (GOF) metric based on the Time Rescaling Theorem (Subramanian & Ramsundar, 2025). If a model correctly captures the heartbeat generative process, the transformed inter-event intervals *z*_*i*_ should be independent and identically distributed uniform random variables on the interval (0, 1).

Let *τ*_*i*_ denote the *i*-th observed RR interval and ℋ_*i*−1_ the history prior to event *i*. If *F* (·|ℋ _*i*−1_) is the model’s conditional cumulative distribution function for the next RR interval, then the time-rescaled variables are

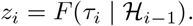

The KS statistic is defined as the maximum deviation between the empirical cumulative distribution of the *z*_*i*_ values and the CDF of a uniform distribution:

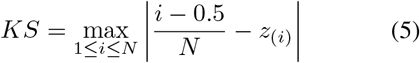

where *z*_(*i*)_ are the sorted transformed intervals. A well-fit model yields a KS score that falls within the 95% significance cutoff, defined as:

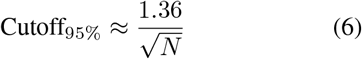

## 3. Experimental Setup and Results

Figure 1 illustrates an example forecast for a particular patient. Qualitatively, we observe that the models cannot capture the long-range stochasticity that is reflected by an RR interval temporal signal, as opposed to the waveform where qualitatively, the TSFMs are capable of emulating.

**Figure 1.**
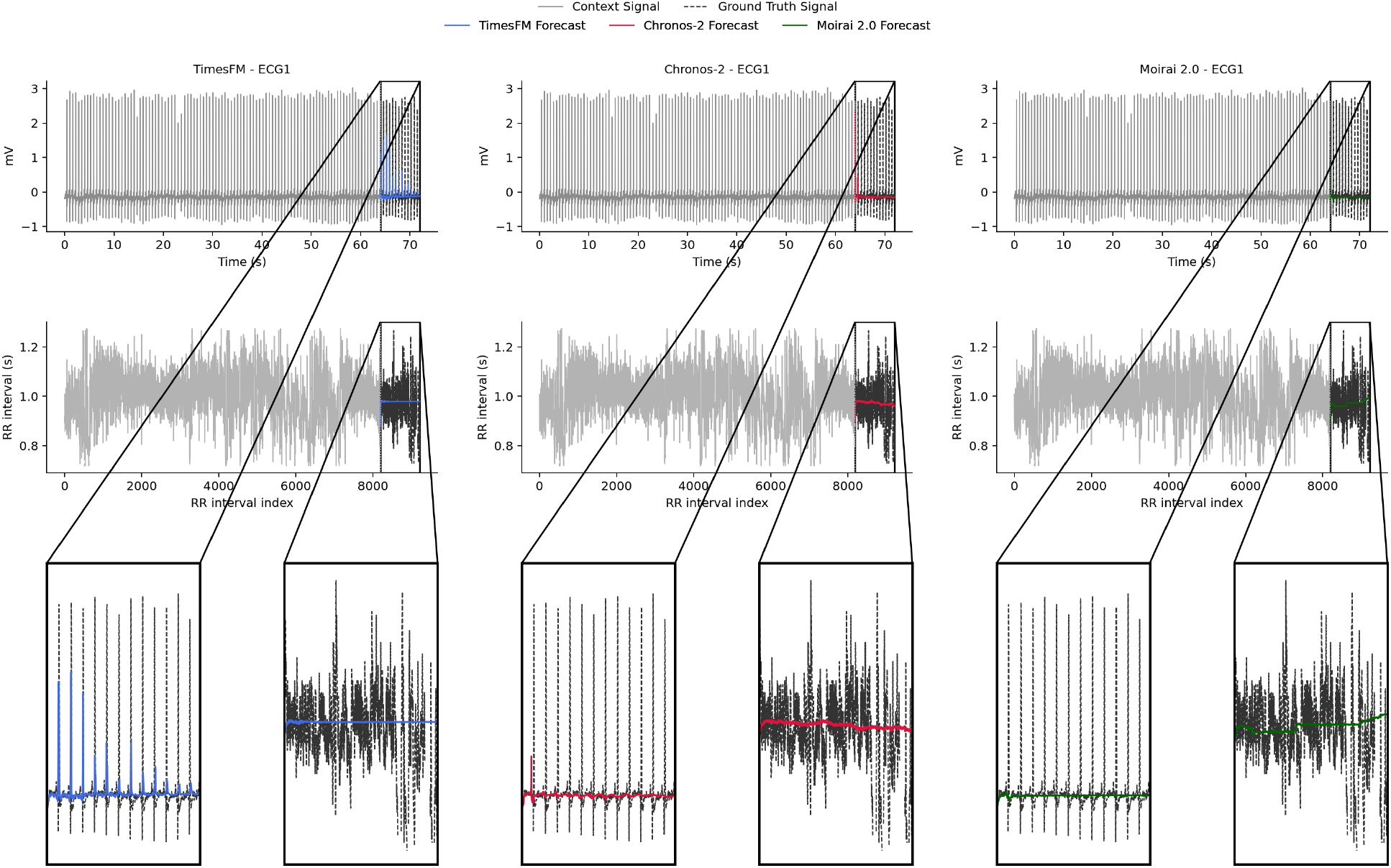
Time-Series Foundation Model (TSFM) forecasts of a single-subject’s ECG data MIT-BIH NSRDB. Each panel compares the observed context window with the forecast horizon for ECG waveform and RR-interval prediction. TSFMs were given 8,192 samples of context and were tasked to forecast 1,024 future waveform or RR interval times.

For our first experiment, we evaluated RR interval forecasting performance across varying ECG context lengths and prediction horizons. This experiment tests whether longer historical context improves forecasting accuracy and whether errors accumulate as models forecast further into the future. We report subject-averaged MAE, RMSE, and KS statistics together with 95% bootstrap confidence intervals across the 18-subject cohort.

As shown in Table 2 (Appendix A), forecasting performance generally worsened as the prediction horizon increased. For example, Chronos-2 direct RR forecasting RMSE increased from 0.17 s (256-beat horizon, 8,192 context) to 0.59 s (1,024-beat horizon, 8,192 context), while TimesFM increased from 0.15 s to 0.34 s over the same horizons, indicating substantial error accumulation with longer forecasts. This degradation was most apparent in RMSE and MAE, particularly for Chronos-2 and Moirai 2.0, suggesting that pointwise prediction errors accumulate over longer horizons. In contrast, increasing the context length from 2,048 to 4,096 or 8,192 samples did not consistently improve performance across models or metrics. The conventional LSTM baseline achieved comparable RMSE and MAE across many settings, indicating that low pointwise forecasting error alone is insufficient to characterize physiologically meaningful long-range forecasting. These results suggest that simply providing additional historical ECG context is insufficient for reliably improving long-range RR interval forecasting.

**Table 2.** Subject-averaged forecasting performance for RR interval prediction across the 18-subject cohort, context lengths, and prediction horizons. Direct denotes direct RR interval prediction; extracted denotes RR intervals derived from model-predicted waveforms via peak extraction. Point estimates are the subject-averaged mean; each is followed by its 95% bootstrap confidence interval (lower, upper) over the 18 subjects.

| Context Length | Prediction Horizon | Output Type | Model | RMSE (s) ↓ | RMSE 95% CI | MAE (s) ↓ | MAE 95% CI | KS ↓ | KS 95% CI |
| --- | --- | --- | --- | --- | --- | --- | --- | --- | --- |
| 2,048 | 256 | Direct | Chronos-2 | 0.24 | (0.17, 0.33) | 0.14 | (0.10, 0.18) | 0.43 | (0.38, 0.47) |
| 2,048 | 256 | Direct | TimesFM | 0.20 | (0.12, 0.29) | 0.11 | (0.07, 0.14) | 0.49 | (0.42, 0.57) |
| 2,048 | 256 | Direct | Moirai 2.0 | 0.49 | (0.44, 0.55) | 0.37 | (0.33, 0.42) | 0.48 | (0.42, 0.53) |
| 2,048 | 256 | Direct | LSTM | 0.18 | (0.10, 0.28) | 0.10 | (0.06, 0.14) | — | — |
| 2,048 | 256 | Extracted | Chronos-2 | 0.14 | (0.11, 0.18) | 0.08 | (0.06, 0.11) | 0.50 | (0.48, 0.52) |
| 2,048 | 256 | Extracted | TimesFM | 0.16 | (0.14, 0.19) | 0.10 | (0.08, 0.12) | 0.51 | (0.49, 0.53) |
| 2,048 | 256 | Extracted | Moirai 2.0 | 0.16 | (0.14, 0.19) | 0.11 | (0.09, 0.14) | 0.44 | (0.43, 0.45) |
| 2,048 | 256 | Extracted | LSTM | 0.24 | (0.19, 0.30) | 0.19 | (0.14, 0.24) | — | — |
| 2,048 | 512 | Direct | Chronos-2 | 0.41 | (0.32, 0.56) | 0.26 | (0.22, 0.30) | 0.46 | (0.43, 0.49) |
| 2,048 | 512 | Direct | TimesFM | 0.23 | (0.12, 0.39) | 0.10 | (0.07, 0.13) | 0.44 | (0.37, 0.52) |
| 2,048 | 512 | Direct | Moirai 2.0 | 0.63 | (0.54, 0.77) | 0.51 | (0.46, 0.56) | 0.61 | (0.57, 0.64) |
| 2,048 | 512 | Direct | LSTM | 0.19 | (0.09, 0.35) | 0.08 | (0.06, 0.10) | — | — |
| 2,048 | 512 | Extracted | Chronos-2 | 0.20 | (0.16, 0.25) | 0.10 | (0.08, 0.13) | 0.46 | (0.44, 0.48) |
| 2,048 | 512 | Extracted | TimesFM | 0.20 | (0.16, 0.24) | 0.10 | (0.08, 0.12) | 0.55 | (0.53, 0.58) |
| 2,048 | 512 | Extracted | Moirai 2.0 | 0.24 | (0.16, 0.34) | 0.12 | (0.09, 0.16) | 0.49 | (0.47, 0.51) |
| 2,048 | 512 | Extracted | LSTM | 0.31 | (0.21, 0.44) | 0.21 | (0.14, 0.30) | — | — |
| 2,048 | 1,024 | Direct | Chronos-2 | 0.66 | (0.50, 0.88) | 0.42 | (0.37, 0.46) | 0.51 | (0.48, 0.54) |
| 2,048 | 1,024 | Direct | TimesFM | 0.34 | (0.15, 0.60) | 0.11 | (0.09, 0.14) | 0.40 | (0.35, 0.45) |
| 2,048 | 1,024 | Direct | Moirai 2.0 | 0.68 | (0.52, 0.90) | 0.44 | (0.38, 0.50) | 0.67 | (0.63, 0.70) |
| 2,048 | 1,024 | Direct | LSTM | 0.32 | (0.13, 0.59) | 0.10 | (0.07, 0.12) | — | — |
| 2,048 | 1,024 | Extracted | Chronos-2 | 0.35 | (0.27, 0.45) | 0.14 | (0.11, 0.16) | 0.44 | (0.42, 0.46) |
| 2,048 | 1,024 | Extracted | TimesFM | 0.25 | (0.20, 0.33) | 0.10 | (0.08, 0.12) | 0.56 | (0.53, 0.58) |
| 2,048 | 1,024 | Extracted | Moirai 2.0 | 0.41 | (0.25, 0.62) | 0.16 | (0.10, 0.23) | 0.52 | (0.50, 0.54) |
| 2,048 | 1,024 | Extracted | LSTM | 0.28 | (0.18, 0.41) | 0.15 | (0.10, 0.19) | — | — |
| 4,096 | 256 | Direct | Chronos-2 | 0.20 | (0.14, 0.27) | 0.10 | (0.08, 0.12) | 0.38 | (0.35, 0.40) |
| 4,096 | 256 | Direct | TimesFM | 0.15 | (0.08, 0.22) | 0.07 | (0.06, 0.09) | 0.46 | (0.39, 0.53) |
| 4,096 | 256 | Direct | Moirai 2.0 | 0.52 | (0.46, 0.58) | 0.41 | (0.36, 0.45) | 0.52 | (0.48, 0.57) |
| 4,096 | 256 | Direct | LSTM | 0.14 | (0.08, 0.22) | 0.06 | (0.05, 0.07) | — | — |
| 4,096 | 256 | Extracted | Chronos-2 | 0.14 | (0.11, 0.17) | 0.08 | (0.06, 0.11) | 0.49 | (0.47, 0.51) |
| 4,096 | 256 | Extracted | TimesFM | 0.16 | (0.14, 0.18) | 0.10 | (0.08, 0.12) | 0.51 | (0.49, 0.53) |
| 4,096 | 256 | Extracted | Moirai 2.0 | 0.15 | (0.13, 0.18) | 0.11 | (0.09, 0.14) | 0.44 | (0.43, 0.45) |
| 4,096 | 256 | Extracted | LSTM | 0.21 | (0.16, 0.26) | 0.16 | (0.11, 0.21) | — | — |
| 4,096 | 512 | Direct | Chronos-2 | 0.36 | (0.27, 0.51) | 0.21 | (0.18, 0.24) | 0.42 | (0.39, 0.45) |
| 4,096 | 512 | Direct | TimesFM | 0.19 | (0.09, 0.35) | 0.08 | (0.06, 0.11) | 0.42 | (0.35, 0.49) |
| 4,096 | 512 | Direct | Moirai 2.0 | 0.64 | (0.55, 0.78) | 0.52 | (0.47, 0.58) | 0.63 | (0.60, 0.67) |
| 4,096 | 512 | Direct | LSTM | 0.19 | (0.09, 0.35) | 0.07 | (0.06, 0.10) | — | — |
| 4,096 | 512 | Extracted | Chronos-2 | 0.23 | (0.17, 0.29) | 0.10 | (0.08, 0.13) | 0.45 | (0.43, 0.47) |
| 4,096 | 512 | Extracted | TimesFM | 0.20 | (0.16, 0.25) | 0.10 | (0.09, 0.12) | 0.55 | (0.53, 0.57) |
| 4,096 | 512 | Extracted | Moirai 2.0 | 0.23 | (0.16, 0.33) | 0.12 | (0.09, 0.15) | 0.49 | (0.46, 0.51) |
| 4,096 | 512 | Extracted | LSTM | 0.24 | (0.17, 0.30) | 0.17 | (0.12, 0.22) | — | — |
| 4,096 | 1,024 | Direct | Chronos-2 | 0.66 | (0.50, 0.89) | 0.41 | (0.37, 0.46) | 0.50 | (0.47, 0.54) |
| 4,096 | 1,024 | Direct | TimesFM | 0.33 | (0.14, 0.60) | 0.10 | (0.08, 0.13) | 0.39 | (0.35, 0.43) |
| 4,096 | 1,024 | Direct | Moirai 2.0 | 0.69 | (0.52, 0.92) | 0.44 | (0.39, 0.51) | 0.67 | (0.64, 0.70) |
| 4,096 | 1,024 | Direct | LSTM | 0.33 | (0.14, 0.61) | 0.10 | (0.08, 0.12) | — | — |
| 4,096 | 1,024 | Extracted | Chronos-2 | 0.39 | (0.31, 0.48) | 0.16 | (0.13, 0.19) | 0.44 | (0.42, 0.46) |
| 4,096 | 1,024 | Extracted | TimesFM | 0.24 | (0.18, 0.30) | 0.10 | (0.08, 0.12) | 0.55 | (0.52, 0.57) |
| 4,096 | 1,024 | Extracted | Moirai 2.0 | 0.34 | (0.22, 0.49) | 0.13 | (0.10, 0.18) | 0.51 | (0.49, 0.53) |
| 4,096 | 1,024 | Extracted | LSTM | 0.28 | (0.21, 0.34) | 0.16 | (0.12, 0.20) | — | — |

Table 2. (continued) Subject-averaged forecasting performance for RR interval prediction across the 18-subject cohort, context lengths, and prediction horizons.
| Context Length | Prediction Horizon | Output Type | Model | RMSE (s) ↓ | RMSE 95% CI | MAE (s) ↓ | MAE 95% CI | KS ↓ | KS 95% CI |
| --- | --- | --- | --- | --- | --- | --- | --- | --- | --- |
| 8,192 | 256 | Direct | Chronos-2 | 0.17 | (0.11, 0.25) | 0.08 | (0.06, 0.10) | 0.33 | (0.30, 0.37) |
| 8,192 | 256 | Direct | TimesFM | 0.15 | (0.09, 0.23) | 0.07 | (0.05, 0.08) | 0.40 | (0.34, 0.47) |
| 8,192 | 256 | Direct | Moirai 2.0 | 0.55 | (0.49, 0.62) | 0.44 | (0.40, 0.49) | 0.56 | (0.51, 0.61) |
| 8,192 | 256 | Direct | LSTM | 0.15 | (0.09, 0.23) | 0.06 | (0.05, 0.08) | — | — |
| 8,192 | 256 | Extracted | Chronos-2 | 0.19 | (0.13, 0.28) | 0.09 | (0.07, 0.12) | 0.47 | (0.45, 0.49) |
| 8,192 | 256 | Extracted | TimesFM | 0.21 | (0.15, 0.28) | 0.10 | (0.08, 0.13) | 0.49 | (0.48, 0.51) |
| 8,192 | 256 | Extracted | Moirai 2.0 | 0.22 | (0.15, 0.30) | 0.13 | (0.10, 0.15) | 0.44 | (0.42, 0.46) |
| 8,192 | 256 | Extracted | LSTM | 0.28 | (0.20, 0.35) | 0.18 | (0.13, 0.23) | — | — |
| 8,192 | 512 | Direct | Chronos-2 | 0.24 | (0.14, 0.40) | 0.12 | (0.09, 0.15) | 0.38 | (0.34, 0.41) |
| 8,192 | 512 | Direct | TimesFM | 0.18 | (0.09, 0.35) | 0.08 | (0.06, 0.11) | 0.39 | (0.33, 0.46) |
| 8,192 | 512 | Direct | Moirai 2.0 | 0.66 | (0.57, 0.80) | 0.55 | (0.51, 0.61) | 0.69 | (0.66, 0.72) |
| 8,192 | 512 | Direct | LSTM | 0.20 | (0.10, 0.36) | 0.09 | (0.06, 0.12) | — | — |
| 8,192 | 512 | Extracted | Chronos-2 | 0.23 | (0.19, 0.28) | 0.11 | (0.09, 0.14) | 0.45 | (0.43, 0.47) |
| 8,192 | 512 | Extracted | TimesFM | 0.20 | (0.16, 0.24) | 0.10 | (0.08, 0.12) | 0.54 | (0.52, 0.56) |
| 8,192 | 512 | Extracted | Moirai 2.0 | 0.18 | (0.14, 0.22) | 0.10 | (0.08, 0.12) | 0.47 | (0.45, 0.50) |
| 8,192 | 512 | Extracted | LSTM | 0.30 | (0.19, 0.44) | 0.20 | (0.12, 0.30) | — | — |
| 8,192 | 1,024 | Direct | Chronos-2 | 0.59 | (0.40, 0.85) | 0.31 | (0.27, 0.35) | 0.47 | (0.43, 0.52) |
| 8,192 | 1,024 | Direct | TimesFM | 0.34 | (0.13, 0.63) | 0.10 | (0.08, 0.12) | 0.37 | (0.34, 0.41) |
| 8,192 | 1,024 | Direct | Moirai 2.0 | 0.76 | (0.58, 1.00) | 0.52 | (0.46, 0.59) | 0.74 | (0.70, 0.77) |
| 8,192 | 1,024 | Direct | LSTM | 0.35 | (0.14, 0.64) | 0.10 | (0.08, 0.13) | — | — |
| 8,192 | 1,024 | Extracted | Chronos-2 | 0.41 | (0.32, 0.50) | 0.16 | (0.13, 0.20) | 0.44 | (0.42, 0.47) |
| 8,192 | 1,024 | Extracted | TimesFM | 0.25 | (0.20, 0.31) | 0.11 | (0.09, 0.12) | 0.54 | (0.52, 0.56) |
| 8,192 | 1,024 | Extracted | Moirai 2.0 | 0.29 | (0.18, 0.43) | 0.12 | (0.08, 0.17) | 0.49 | (0.47, 0.52) |
| 8,192 | 1,024 | Extracted | LSTM | 0.35 | (0.18, 0.65) | 0.18 | (0.11, 0.28) | — | — |

For our next experiment, we compared two approaches for RR interval forecasting: direct prediction and extraction from model-generated ECG waveforms. Extracted RR intervals often achieved lower RMSE and MAE, indicating better pointwise accuracy. However, this advantage did not consistently extend to the KS metric. In several cases, direct prediction yielded lower KS despite higher RMSE/MAE, while in others the reverse held, suggesting a tradeoff between pointwise accuracy and distributional fidelity.

Lastly, we further analyzed error as a function of forecast beat index to characterize temporal degradation. As shown in Figure 2, all models exhibit increasing error, but with distinct dynamics. TimesFM shows gradual error growth, whereas Moirai 2.0 degrades rapidly and saturates early, and Chronos-2 exhibits delayed but sharp escalation at longer horizons. Critically, both TimesFM and Moirai 2.0 collapse in local variability (SDNN, RMSSD), producing near-zero RR variability relative to the ground truth. Moreover, all three models fail to adequately capture physiological variability of the ECG signal. This indicates that despite reasonable short-horizon accuracy, these models fail to preserve physiologically meaningful dynamics.

**Figure 2.**
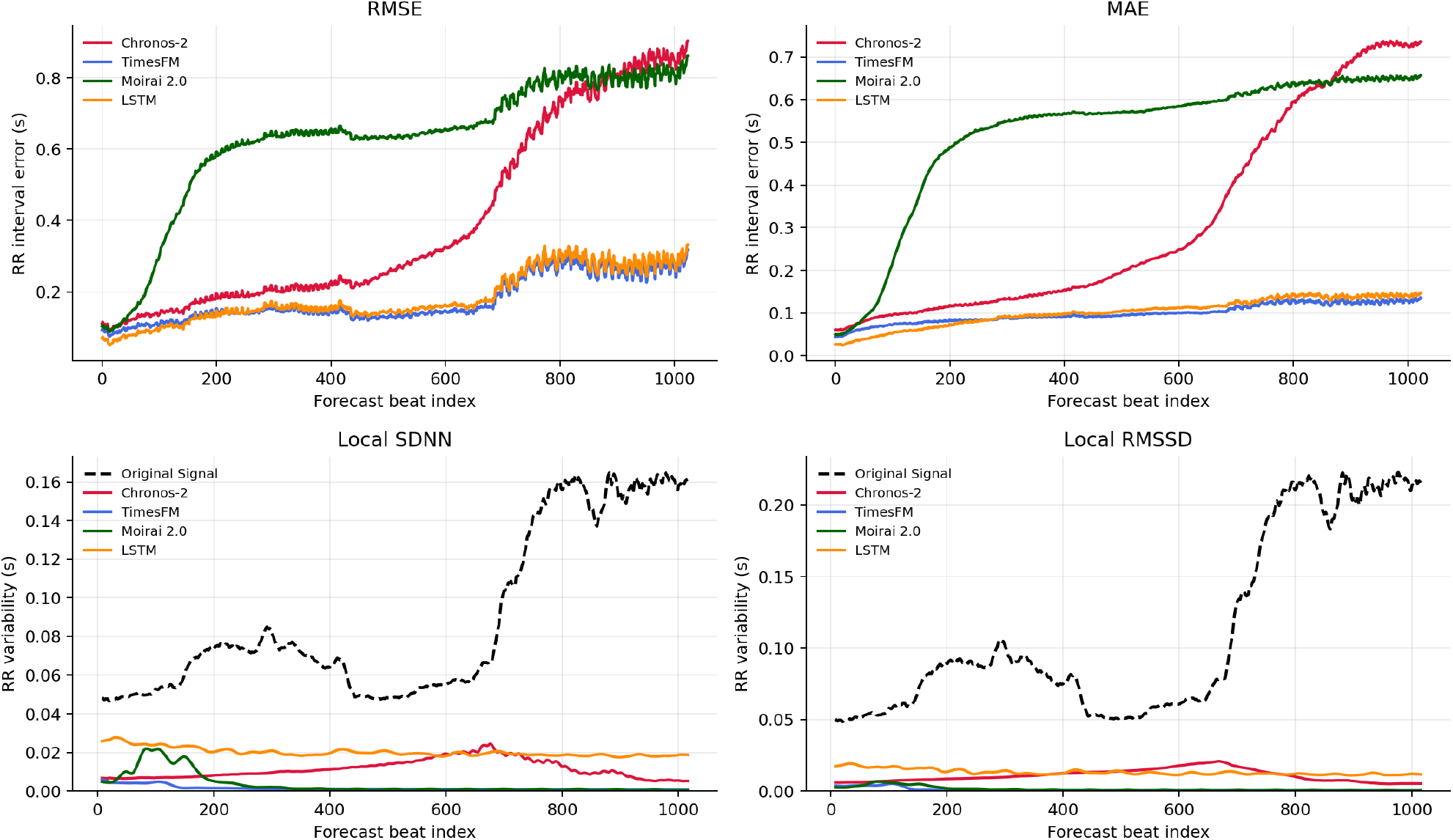
Per-beat evaluation of RR interval forecasting error and variability, averaged across the 18-subject cohort, for forecasts conditioned on the longest evaluated context window (8,192 samples). The RMSE and MAE are plotted as a function of forecast beat index. Local variability metrics (SDNN and RMSSD) are compared to the ground truth RR interval signal. (Note: Table 2 evaluates the model at different context windows, while Figure 2 evaluates at a single context window)

## 4. Discussion

The results highlight both the promise and limitations of Time-Series Foundation Models (TSFMs) for physiological data. While these models generate realistic short-term ECG waveforms, TSFM forecasting capabilities does not necessarily extend to long-range temporal scales. Analysis of RR interval dynamics reveals that performance degrades when modeling long-range structure.

A central question is whether TSFMs capture physiologically meaningful dynamics or primarily reproduce local statistical patterns. We observe that the models appear to rely on beat-level morphology rather than learning long-range cardiac behavior. This distinction is critical, as many clinically relevant processes such as autonomic regulation emerge over longer time scales. RR intervals provide a more direct probe of this limitation. Unlike waveform morphology, which can be matched locally, RR interval structure reflects integrated physiological control. Failure to model these dynamics indicates limited ability to capture meaningful temporal hierarchies.

These results have important implications for clinical deployment. Although TSFMs are attractive in low-data settings, caution is warranted for tasks requiring long-range physiological consistency.

Future work should prioritize evaluation frameworks that explicitly test long-range temporal modeling, such as RR interval forecasting. Moreover, other metric such as the Fréchet distance (Tang et al., 2025) could further evaluate the statistical validity and physiological fidelity of model forecasts. Improving performance may require architectural biases, extended context windows, and training objectives that enforce multi-scale temporal structure in physiological signals.

## A. RR Forecast Context Length and Prediction Horizon Experiment

